# Tamoxifen repurposing to combat infections by multidrug-resistant Gram-negative bacilli

**DOI:** 10.1101/2020.03.30.017475

**Authors:** Andrea Miró-Canturri, Rafael Ayerbe-Algaba, Raquel del Toro, Jerónimo Pachón, Younes Smani

**Affiliations:** Clinical Unit of Infectious Diseases, Microbiology and Preventive Medicine, University Hospital Virgen del Rocío, Seville, Spain; Institute of Biomedicine of Seville (IBiS), University Hospital Virgen del Rocío/CSIC/University of Seville, Seville, Spain; Department of Medical Physiology and Biophysics, Institute of Biomedicine of Seville, University of Seville, Sevilla, Spain; Department of Medicine, University of Seville, Seville, Spain; CIBER de enfermedades cardiovasculares (CIBER-CV), Spain

**Author notes:** To whom correspondence should be addressed: Younes Smani, Clinic Unit of Infectious Diseases, Microbiology and Preventive Medicine, Institute of Biomedicine of Seville (IBiS), University Hospital Virgen del Rocío, Av. Manuel Siurot s/n, 41013, Seville, Spain. Both authors have contributed equally to this work (Andrea Miró-Canturri, Rafael Ayerbe-Algaba).

## Abstract

The development of new strategic therapies for multidrug-resistant bacteria, like the use of non-antimicrobial approaches and/or drugs repurposing to be used as monotherapies or in combination with clinically relevant antibiotics, has become an urgent need. A therapeutic alternative for infections by multidrug-resistant Gram-negative bacilli (MDR-GNB) is immune system modulation to improve the infection clearance. We showed that immunocompetent mice infected by *Acinetobacter baumannii, Pseudomonas aeruginosa* or *Escherichia coli* in peritoneal sepsis models and treated with tamoxifen at 80 mg/kg/d for three days reduced the release of MCP-1 and its signalling pathway IL-18 and phosphorylated ERK1/2. This reduction of MCP-1 induced the reduction of migration of inflammatory monocytes and neutrophils from bone marrow to blood. Indeed, the treatment with tamoxifen in murine peritoneal sepsis models reduced the bacterial load in tissues and blood; and increased the mice survival from 0% to 60-100%. Tamoxifen treatment of neutropenic mice infected by these pathogens increased mice survival up to 20-60%. Furthermore, susceptibility and time-kill assays showed that the metabolites of tamoxifen, N-desmethyltamoxifen, hydroxytamoxifen and endoxifen, the three together exhibited MIC_90_ values of 16 mg/L and were bactericidal against clinical isolates of *A. baumannii* and *E. coli*. This antimicrobial activity of tamoxifen metabolites parallels’ an increased membrane permeability of *A. baumannii* and *E. coli* without affecting their outer membrane proteins profiles. Together, these data showed that tamoxifen present a therapeutic efficacy against MDR *A. baumannii, P. aeruginosa* and *E. coli* in experimental models of infections and can be repurposed as new treatment for GNB infections.

**Importance:** Antimicrobial resistance in Gram-negative bacilli (GNB) is a global health treat. Drug repurposing, a novel approach involving the search of new indications for FDA approved drugs is gaining interest. Among them, we found the anti-cancer drug tamoxifen, which presents very promising therapeutic efficacy. The current study showed that tamoxifen presents activity in animal models of infection with MDR *Acinetobacter baumannii, Pseudomonas aeruginosa* and *Escherichia coli* by modulating the traffic of innate immune system cells and the antibacterial activity presented by its three major metabolites produced *in vivo* against these GNB. Our results offer a new candidate to be repurposed to treat severe infections caused by these pathogens.

## Introduction

Infections caused by Gram-negative bacilli (GNB) such as *Acinetobacter baumannii, Pseudomonas aeruginosa* and *Escherichia coli* represent an increasing worldwide problem. In 2017, the World Health Organization has listed these pathogens as the first antibiotic-resistant “priority pathogens” that pose the greatest threat to human health. There is, therefore, an urgent need to find new antimicrobial agents against extensive- and pan-drug-resistant GNB. Two key approaches can help alleviate the problem of antibiotic resistance, firstly targeting of bacterial virulence factors without inhibiting bacterial growth, which can slow the development of drug resistance by reducing the selective pressure on the bacteria (1, 2) and, secondly, by the modulation/regulation of the immune system response to improve the infection development (3, 4). In this way, some studies were focused on the stimulation of the immune system to treat bacterial infections using molecules, including lysophosphatidylcoline as monotherapy and as adjuvant for the antimicrobial treatment (3, 5, 6) or 3’-5’-cyclic diguanylic acid (c-di-GMP) which increase neutrophils protecting against *A. baumannii* infection (7).

Inflammatory monocytes and neutrophils derived from bone marrow are important cellular mediators of innate immune response against bacterial infections. During early stages of bacterial infection, both cell populations migrate from the bone marrow to the bloodstream and subsequently to the sites of infection (8, 9). This migration is regulated partially by the monocyte chemotactic protein-1 (MCP-1), which expression is increased by bone marrow mesenchymal cells in response to circulating Toll-like receptor ligands and produces the mobilization of inflammatory monocytes (10). It is well established that MCP-1 release is controlled by IL-18 and ERK1/2 (11), and the levels of MCP-1 are higher in patients with sepsis and septic shock, and pneumonia (12, 13).

It is well documented that anti-cancer drugs like tamoxifen can modify the immune response by regulating cytokine release (14). Mechanistically, tamoxifen has been reported to reduce MCP-1 transcription and expression in human coronary artery endothelial cells and endometrial cancer cells, respectively (15, 16). As MCP-1 is involved in the immune cells migration, it may be hypothesized that an undiscovered connection between MCP-1 release and immune cells migration after bacterial infection and treatment with tamoxifen is present.

In prokaryotic cells, tamoxifen is known to present antifungal and antibacterial activities against *Mycobacterium tuberculosis* and some Gram-positive bacteria *in vitro* and *in vivo* (17, 18), but not against Gram-negative bacteria. As with other antimicrobial agent such as colistimethate sodium (19), tamoxifen is a prodrug and converted after liver passage to three major active metabolites, 4-hydroxytamoxifen, endoxifen and N-desmethyltamoxifen (20). However, their antibacterial activities remain unknown.

In this study, we report that tamoxifen downregulates the expression of MCP-1, impairing the migration of bone marrow derived cells to the bloodstream induced by *A. baumannii, P. aeruginosa* and *E. coli* and, consequently, modulating the inflammatory response. In murine peritoneal sepsis model, we observe that tamoxifen decreases the development of infection by these pathogens, lowering their concentrations in tissues and blood and increasing the mice survival. Although tamoxifen did not present bactericidal nor bacteriostatic effects against *A. baumannii, P. aeruginosa* and *E. coli in vitro*, we show that tamoxifen metabolites exhibit high antibacterial activity against *A. baumannii* and *E. coli*, suggesting that tamoxifen metabolism is actively involved in the therapeutic efficacy of tamoxifen *in vivo*.

## Results

### Bone marrow immune cells migrates in response to MCP-1 and IL-18 during bacterial infection

To determine whether bacterial infection influences circulating immune cells from the bone marrow in response to MCP-1 and IL-18, a MCP-1 controller (11), we administered intraperitoneally *A. baumannii, P. aeruginosa* and *E. coli* to mice and measured the proportions of myeloid cells CD11b+, inflammatory monocytes CD11b+Ly6C^hi^ and neutrophils CD11b+Ly6G+. *A. baumannii* administration decreased, 24 h after, the myeloid cells, inflammatory monocytes and neutrophils in bone marrow, and increased them in blood (Figures 1A, 1B and 1C). Same results were observed when mice were infected with *P. aeruginosa* and *E. coli* (Figures 1A, 1B and 1C). The rates of these immune cells in the spleen remained unchanged after infection with *A. baumannii, P. aeruginosa* and *E. coli* for 24 h (Figure S1) indicating that the increase of circulating monocytes and neutrophils did not proceed from the splenic reservoir (21, 22).

**Figure 1.**
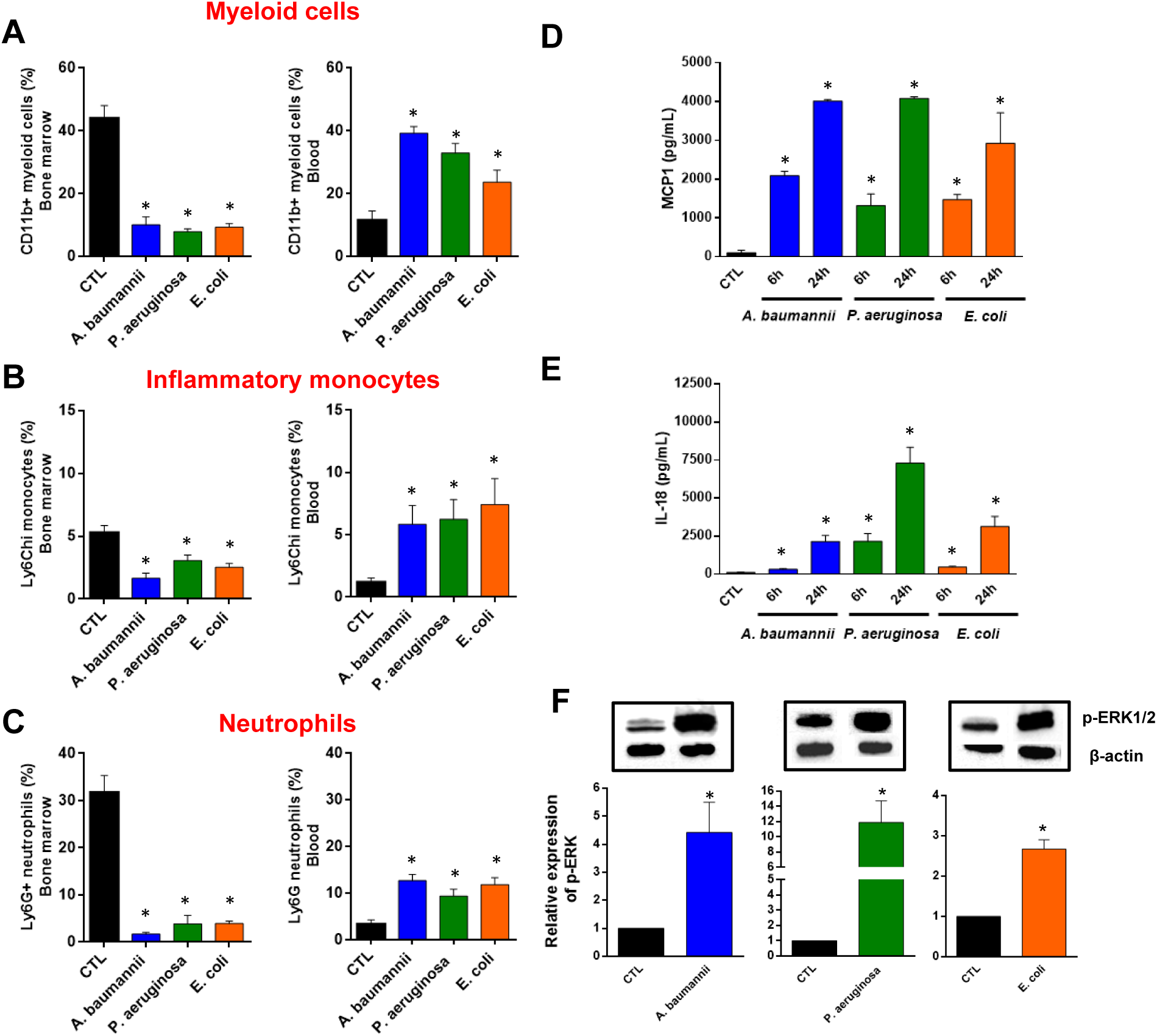
Bone marrow immune cells migration to blood in response to MCP-1 and IL-18 during bacterial infection. (**A**) Myeloid cells, (**B**) inflammatory monocytes and (**C**) neutrophils were identified as CD11b+, CD11b+Ly6Chi and CD11b+Ly6G+ by flow cytometry, respectively, in bone marrow and blood of mice infected with MLD100 of *A. baumannii* ATCC17978, *P. aeruginosa* PAO1 or *E. coli* ATCC25922 strains for 24h. (**D** and **E**) Serum MCP-1 and IL-18 levels (ELISA assays), 6 and 24 h post-infection, in mice infected with minimal lethal dose 100 (MLD100) of *A. baumannii* ATCC17978, *P. aeruginosa* PAO1 or *E. coli* ATCC25922 strains. (**F**) RAW 264.7 cells were infected with *A. baumannii* ATCC17978, *P. aeruginosa* PAO1 or *E. coli* ATCC25922 strains for 2 h and proteins were collected for Phospho-p44/42 MAPK (Erk1/2) and β-actin immunoblotting. Data are representative of six mice per group, and expressed as mean ± SEM. \**P*<0.05: infected vs. CTL. CTL: non-infected mice.

A paradigm widely accepted is the formation of chemokine gradients to guide inflammatory cells to the sites of infection (23). Among them, MCP-1 has been shown to be involved in the migration of immune cells from the bone marrow to the bloodstream after binding to CCR2 receptor (24). As it is shown in the figure 1D, mice infected with *A. baumannii, P. aeruginosa* and *E. coli* for 6 and 24 h increased significantly and progressively the release of MCP-1 in mice serum (between ≈1000 and 4000 µg/mL). It is well known that MCP-1 release is controlled by IL-18 and ERK1/2 (11). Consequently, the levels of IL-18 in mice serum gradually increased 6 and 24 h after infection by *A. baumannii, P. aeruginosa* and *E*.*coli*. The IL-18 levels at 24 h were 2144 ± 408.1 µg/mL, 7286 ± 1056 µg/mL and 3124 ± 671.3 µg/mL, respectively (Figure 1E). Moreover, ERK1/2 was phosphorylated 2 h after infection of RAW 264.7 macrophages cell line *in vitro* by *A. baumannii, P. aeruginosa* and *E. coli*, defining the activation of kinase response to these pathogens (Figure 1F).

To determine whether MCP-1 is involved in the migration of inflammatory monocytes and neutrophils from bone marrow to blood, wild-type (WT) and mice lacking MCP-1 protein (MCP-1 KO mice) were infected by *A. baumannii, P. aeruginosa* and *E. coli*. First, we detected MCP-1 release only in WT mice (Figure 2A). Importantly, the infection of MCP-1 KO mice by these pathogens showed that the migration of inflammatory monocytes and neutrophils from bone marrow to blood (Figures 2B and 2C) exhibits a reduction of 2.17 ± 1.14% and 4.13 ± 0.99%, respectively, for *A. baumannii* infection. Similar results were observed when MCP-1 KO mice were infected with *P. aeruginosa* and *E. coli* strains (Figures 2B and 2C).

**Figure 2.**
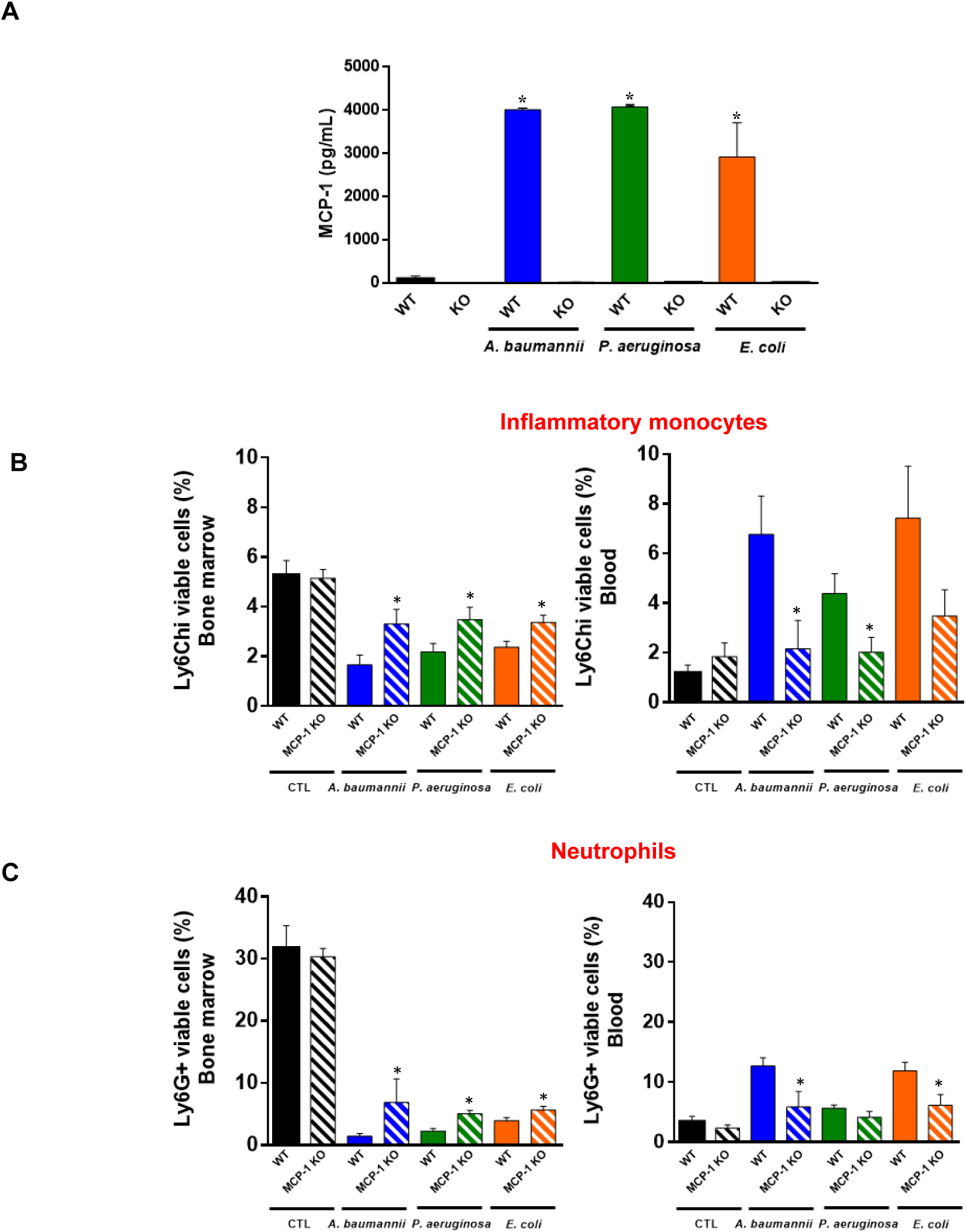
Role of MCP-1 in the bone marrow immune cells migration to blood during bacterial infection. (**A**) Wild-type and MCP-1 KO mice were infected with minimal lethal dose 100 (MLD100) of *A. baumannii* ATCC17978, *P. aeruginosa* PAO1 or *E. coli* ATCC25922 strains. Twenty-four hours post-infection, serum was harvested for MCP-1 ELISA assays. (**B**) Inflammatory monocytes, and (**C**) neutrophils were identified as CD11b+Ly6Chi and CD11b+Ly6G+ by flow cytometry, respectively, in bone marrow and blood of wild-type and MCP-1 KO mice infected with MLD100 of *A. baumannii* ATCC17978, *P. aeruginosa* PAO1 or *E. coli* ATCC25922 strains for 24 h. Data are representative of six mice per group, and expressed as mean ± SEM. \**P*<0.05: WT *vs*. MCP-1 KO. WT: wild-type, MCP-1 KO: mice lacking MCP-1, CTL: non-infected mice.

Non-infected WT and MCP-1 KO mice presented similar inflammatory monocytes and neutrophil proportions in bone marrow and blood indicating that the lack of MCP-1 did not affect the migration of these cells from bone marrow in basal conditions (Figures 2B and 2C). These data suggest that MCP-1 is involved in the traffic of immune cells from the bone marrow to blood after infection by *A. baumannii, P. aeruginosa* and *E. coli*.

### Tamoxifen impairs the migration of immune cells from bone marrow to blood through MCP-1 regulation

In order to study whether tamoxifen can modulate inflammation generated by bacterial infections, we treated RAW 264.7 macrophage cell line with tamoxifen during 24 h and infected them with *A. baumannii, P. aeruginosa* or *E. coli* for 2 h. After this incubation we determined the secretion of MCP-1 in the macrophage cells supernatant (ELISA assay) and the phosphorylation of ERK in the macrophage cells by Western blot. The treatment with tamoxifen decreased the release of MCP-1 and the phosphorylation of ERK1/2 in macrophages infected by these pathogens, compared to macrophages without tamoxifen treatment (Figures 3A and 3B). To confirm these data *in vivo*, mice were treated ip. with 3 doses of 80 mg/kg/d of tamoxifen before the bacterial infection. Serum was collected 6 and 24 h post-bacterial infection. Figure 3C revealed that treatment with tamoxifen reduced MCP-1 levels when compared with *A. baumannii, P. aeruginosa* or *E. coli* infected and not treated groups. It is noteworthy to highlight that IL-18 levels were also reduced after tamoxifen treatment of infected mice by *A. baumannii, P. aeruginosa* and *E. coli* (Figure 3D). These results suggest that the reduction of IL-18 secretion due to tamoxifen injection may drive a reduction of MCP-1 release through a reduction of ERK phosphorylation. This MCP-1 reduction after tamoxifen injection could produces less migration of monocytes and neutrophils from the bone marrow to the blood.

**Figure 3.**
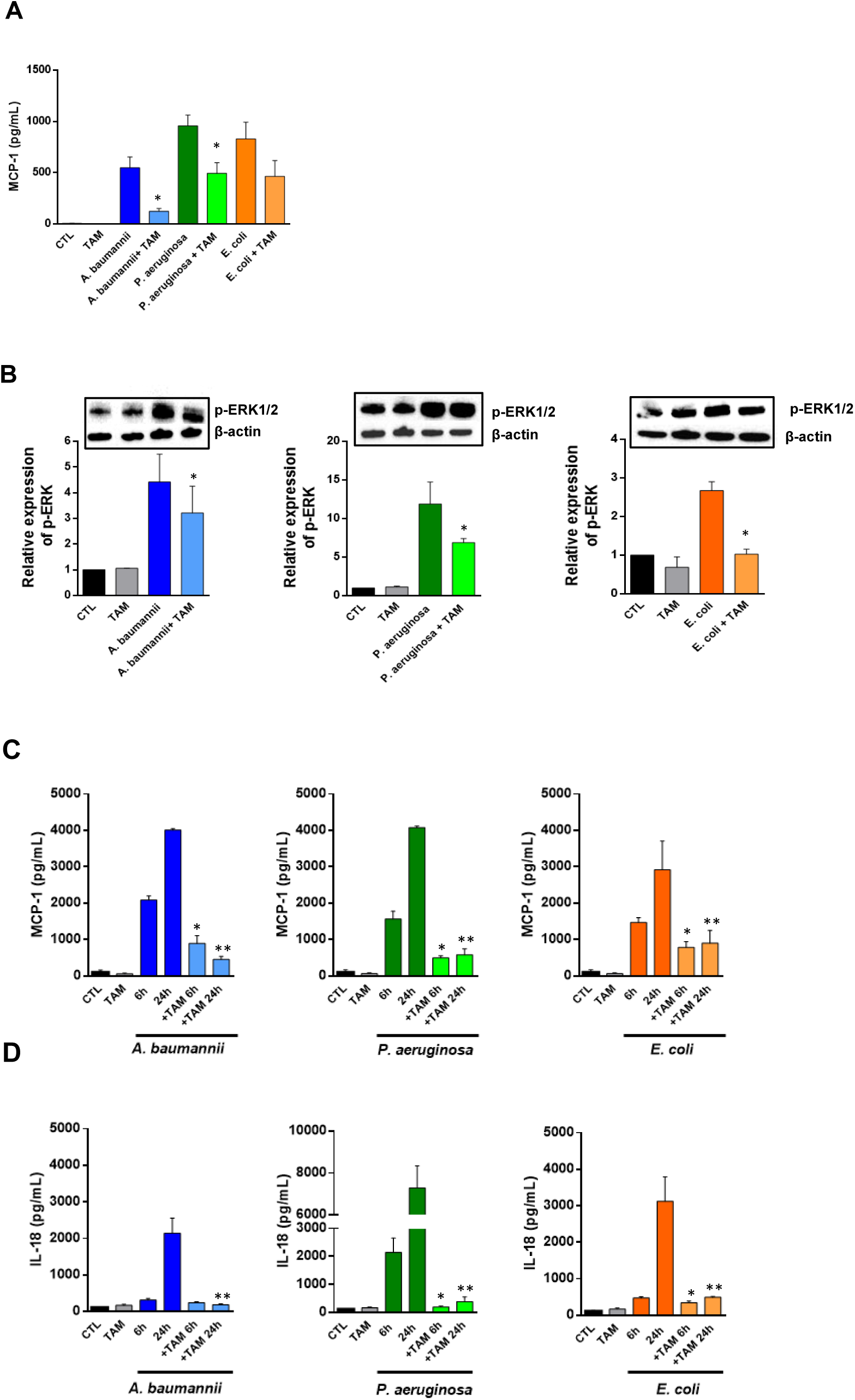
Tamoxifen reduces, after bacterial infection, the release of MCP-1 and IL-18 *in vitro* and *in vivo*, and the ERK phosphorylation *in vitro*. (**A** and **B**) RAW 264.7 cells were treated with 2.5 mg/L of tamoxifen for 24 h and infected with *A. baumannii* ATCC17978, *P. aeruginosa* PAO1 or *E. coli* ATCC25922 strains for 2 h. MCP-1 levels and ERK-phosphrylation were determined by ELISA and immunoblotting assays, respectively. Data are representative of three independent experiments, and expressed as mean ± SEM. (**C** and **D**) Mice received tamoxifen (80 mg/kg/d, for 3 days) and infected with minimal lethal dose 100 (MLD100) of *A. baumannii* ATCC17978, *P. aeruginosa* PAO1 or *E. coli* ATCC25922 strains. Six and twenty-four hours post-infection, serum was harvested for MCP-1 and IL-18 ELISA assays. Data are representative of 6 mice per group and are expressed as mean ± SEM. \**P*<0.05: treated *vs*. CTL, \*\**P*<0.05: treated *vs*. CTL. CTL: non-infected mice. TAM: tamoxifen.

In order to confirm whether tamoxifen treatment reduces the proportions of myeloid cells, inflammatory monocytes and neutrophils in bone marrow and blood, we administer tamoxifen in mice before infection by *A. baumannii, P. aeruginosa* and *E. coli* for 24 h. Flow cytometric analysis demonstrated that treatment with tamoxifen reduced the migration of these cells to the blood and the levels in bone marrow were maintained compared with the levels of infected group (Figures 4A, 4B and 4C).

**Figure 4.**
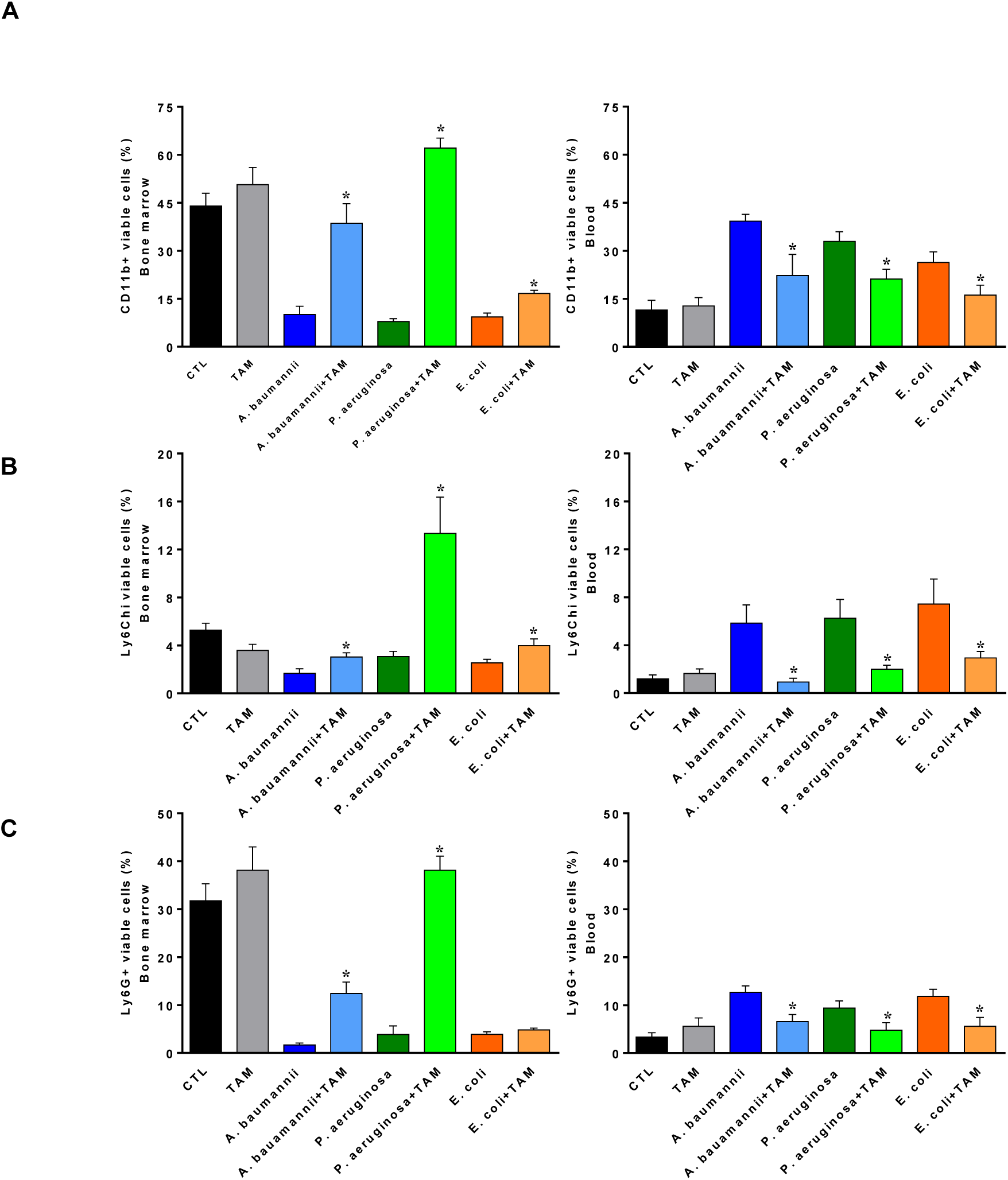
Tamoxifen impairs, after bacterial infection, the migration of immune cells from bone marrow to blood through MCP-1 regulation. Mice received tamoxifen (80 mg/kg/d, for 3 days) and infected with minimal lethal dose 100 of *A. baumannii* ATCC17978, *P. aeruginosa* PAO1 or *E. coli* ATCC25922 strains. Twenty-four hours post-infection, (**A**) myeloid cells, (**B**) inflammatory monocytes and (**C**) neutrophils were identified as CD11b+, CD11b+Ly6Chi and CD11b+Ly6G+ by flow cytometry, respectively, in bone marrow and blood of mice. Data are representative of 6 mice per group and are expressed as mean ± SEM. \**P*<0.05: treated *vs*. CTL. CTL: non-infected mice, TAM: tamoxifen.

MCP-1 KO mice showed an impaired migration of inflammatory monocytes and neutrophils from bone marrow to blood after bacterial infection (Figures 2B and 2C). In order to determine whether tamoxifen is able to reduce this migration in mice deficient in MCP-1 secretion, we treated MCP-1 KO mice with tamoxifen and infected them with *A. baumannii, P. aeruginosa* and *E. coli*. As it is showed in the figures 5A and 5B, tamoxifen treated mice presented a reduction in the migration of inflammatory monocytes and neutrophils, despite of the lack of MCP-1. Both populations were more present in bone marrow and the frequencies in the blood were also reduced when compared with WT mice treated with tamoxifen and infected by these pathogens, indicating that tamoxifen may probably regulate other chemokines and migration pathways involved in this phenomenon (Figure S2, Figure 5).

**Figure 5.**
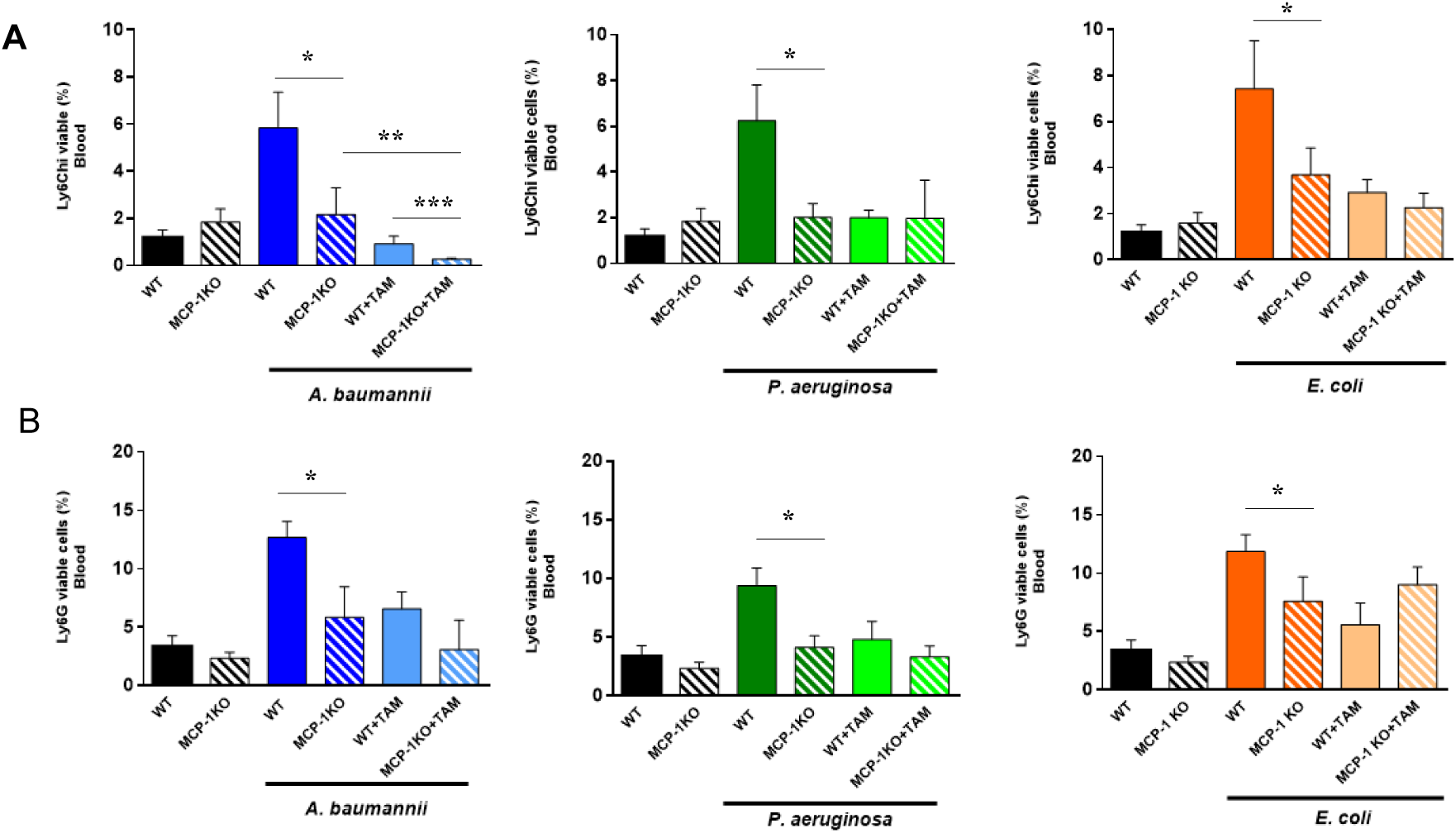
Tamoxifen impairs, after bacterial infection in mice MCP-1-deficient, the migration of immune cells from bone marrow to blood through MCP-1 regulation. (**A**) WT and MCP-1 KO mice received tamoxifen (80 mg/kg/d, for 3 days) and infected with minimal lethal dose 100 of *A. baumannii* ATCC17978, *P. aeruginosa* PAO1 or *E. coli* ATCC25922 strains. Twenty-four hours post-infection, (**A**) inflammatory monocytes and (**B**) neutrophils were identified as CD11b+, CD11b+Ly6Chi and CD11b+Ly6G+ by flow cytometry, respectively, in bone marrow and blood of mice. Data are representative of 6 mice per group and are expressed as mean ± SEM. CTL: non-infected mice. \**P*<0.05: infected WT *vs*. infected MCP-1 KO, \*\**P*<0.05: infected MCP-1 KO *vs*. infected MCP-1 KO + TAM, \*\*\**P*<0.05: infected WT + TAM *vs*. infected MCP-1 KO + TAM. CTL: non-infected mice, TAM: tamoxifen.

### Tamoxifen enhances bacterial killing of macrophages and neutrophils in-vitro

Recent studies reported that treatment with tamoxifen enhances neutrophil activity by increasing the NETosis and induces changes in macrophages by inhibiting the expression of CD36 and PPARγ reducing atherosclerosis (25, 26), but there are no data regarding the immune function of both cells treated with tamoxifen after a bacterial infection. To determine whether tamoxifen can increase the killing activity of macrophages and neutrophils, assays with RAW 246.7 cell line and HL-60 neutrophils cell line pretreated with tamoxifen and infected by *A. baumannii, P. aeruginosa* and *E. coli* were performed. We demonstrated that macrophages incubation with tamoxifen (2.5 mg/L) at 2 and 6 h followed by infection with *A. baumannii* during 2 h decreased the bacterial internalization by 10 and 30%, respectively (Figure S3A), without affecting the amount of *A. baumannii* in the extracellular medium (Figure S3B). Similar results were observed after treatment with tamoxifen and infection by *E. coli*, but not by *P. aeruginosa* (Figure S3A). Regarding neutrophil activity, incubation with 2.5 mg/L of tamoxifen during 2 and 6 h followed by the infection with *A. baumannii* during 2 h increased bacterial killing by 5 and 25%, respectively. Similar results were observed after treatment with tamoxifen and infection by *E. coli*, but not by *P. aeruginosa* (Figure S3A).

Accordingly, tamoxifen treatment increases the killing activity of macrophages and neutrophils against *A. baumannii* and *E. coli* but not against *P. aeruginosa*.

### Tamoxifen increase mice survival and decrease the bacterial burden in a murine sepsis model by A. baumannii, P. aeruginosa and E. coli

Our results demonstrated that tamoxifen plays an important role in innate immune cells trafficking after bacterial infection. Going further we wanted to know whether tamoxifen could protect the mice against a lethal bacterial inoculum. We treated mice with tamoxifen (80 mg/kg/d) administered intraperitoneally three days before the infection with a minimal lethal dose 100 (MLD100) of *A. baumannii, P. aeruginosa* and *E. coli*. Pretreatment with tamoxifen increased mice survival after infection by *A. baumannii, P. aeruginosa* and *E. coli* to 100, 66.7 and 83.3% (*P*<0.01), respectively (Figure 6A). Figure 6B shows that treatment with tamoxifen decreased spleen and lung bacterial concentrations of these pathogens by 6.64 and 7.17 log_10_ CFU/g (*P*<0.015; for *A. baumannii*), by 3.58 and 5.1 log_10_ CFU/g (*P*<0.015; for *P. aeruginosa*), and by 3.7 and 4.16 log_10_ CFU/g (*P*<0.015; for *E. coli*), compared with the control infected groups. Blood bacterial concentrations presented a decrease compared to control infected groups of 5.53, 5.45 and 4.31 log_10_ CFU/mL (*P*<0.01) for *A. baumannii, P. aeruginosa* and *E. coli*, respectively. Similar efficacy of tamoxifen was observed in murine peritoneal sepsis model by susceptible and MDR clinical isolates of *A. baumannii, P. aeruginosa* and *E. coli*. Treatment with tamoxifen increased the mice survival to 66.7, 83.3 and 50% (*P*<0.01) for the non-MDR *A. baumannii, P. aeruginosa* and *E. coli*, respectively, and 83.3, 66.7 and 50% (*P*<0.01) for the MDR *A. baumannii, P. aeruginosa* and *E. coli* harboring *mcr-1* gene (Figure 6C). These findings indicate that tamoxifen treatment presents a good therapeutic efficacy against reference and clinical isolates of *A. baumannii, P. aeruginosa* and *E. coli*.

**Figure 6.**
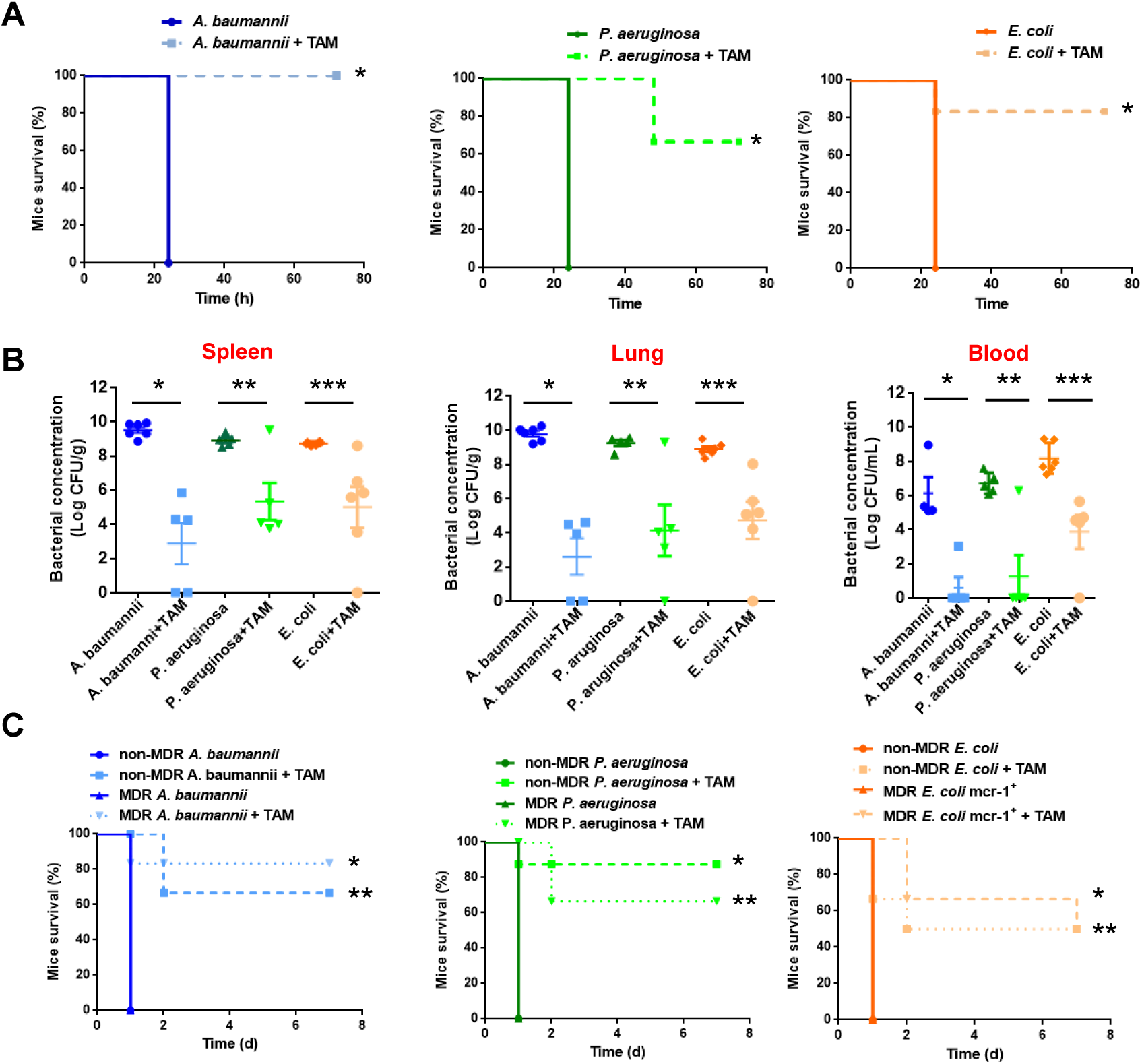
Tamoxifen shows therapeutic efficacy in murine sepsis models by GNB. (**A**) Mice survival was monitored during 3 days for mice infected with minimal lethal dose 100 (MLD100) of *A. baumannii* ATCC17978, *P. aeruginosa* PAO1 or *E. coli* ATCC25922 strains treated or not with 3 i.p. doses of tamoxifen (80 mg/kg/d, for 3 days). (**B**) Bacterial burden in spleen, lung and blood of mice treated or not with 3 ip. doses of tamoxifen (80 mg/kg/d, for 3 days), and infected with MLD100 of *A. baumannii* ATCC17978, *P. aeruginosa* PAO1 or *E. coli* ATCC25922 strains. (**C**) Mice survival was monitored during 7 days for 6 mice infected with MLD100 of non-MDR and MDR *A. baumannii* (Ab9 and Ab186), *P. aeruginosa* (Pa39 and Pa238) or *E. coli* (C1-7-LE and EcMCR+) strains treated or not with 3 i.p. doses of tamoxifen (80 mg/kg/d, for 3 days). \**P*<0.05: treated *vs*. untreated, \*\**P*<0.05: treated *vs*. untreated, \*\*\**P*<0.05: treated *vs*. untreated. TAM: tamoxifen, MDR: multidrug-resistant.

### Immunosuppressed mice respond to tamoxifen treatment

Previous studies have demonstrated that infection with *A. baumannii, P. aeruginosa* and *E. coli* in immunosuppressed mice is lethal (27-29). To determine whether tamoxifen treatment is still therapeutically effective in immunosuppressed mice, we treated immunocompetent mice with cyclophosphamide to reduce the circulating monocytes and neutrophils (Figure S4). After *A. baumannii* and *E. coli* infection in these immunosuppressed mice, tamoxifen treatment increase mice survival in both groups to 66.67 % (Figure 7A); however, with *P. aeruginosa* the survival was only 16.67% (Figure 7A). Bacterial loads of *A. baumannii* and *E. coli* in spleen, lung and blood were reduced in immunosuppressed mice after treatment with tamoxifen, likewise in immunocompetent mice. In contrast, bacterial loads of *P. aeruginosa* in tissues and blood were not reduced in immunosuppressed mice after treatment with tamoxifen (Figure 7B). These findings suggest that tamoxifen help to clear the infection by *A. baumannii* and *E. coli* even though mice were immunosuppressed by an additional independent immune response mechanism.

**Figure 7.**
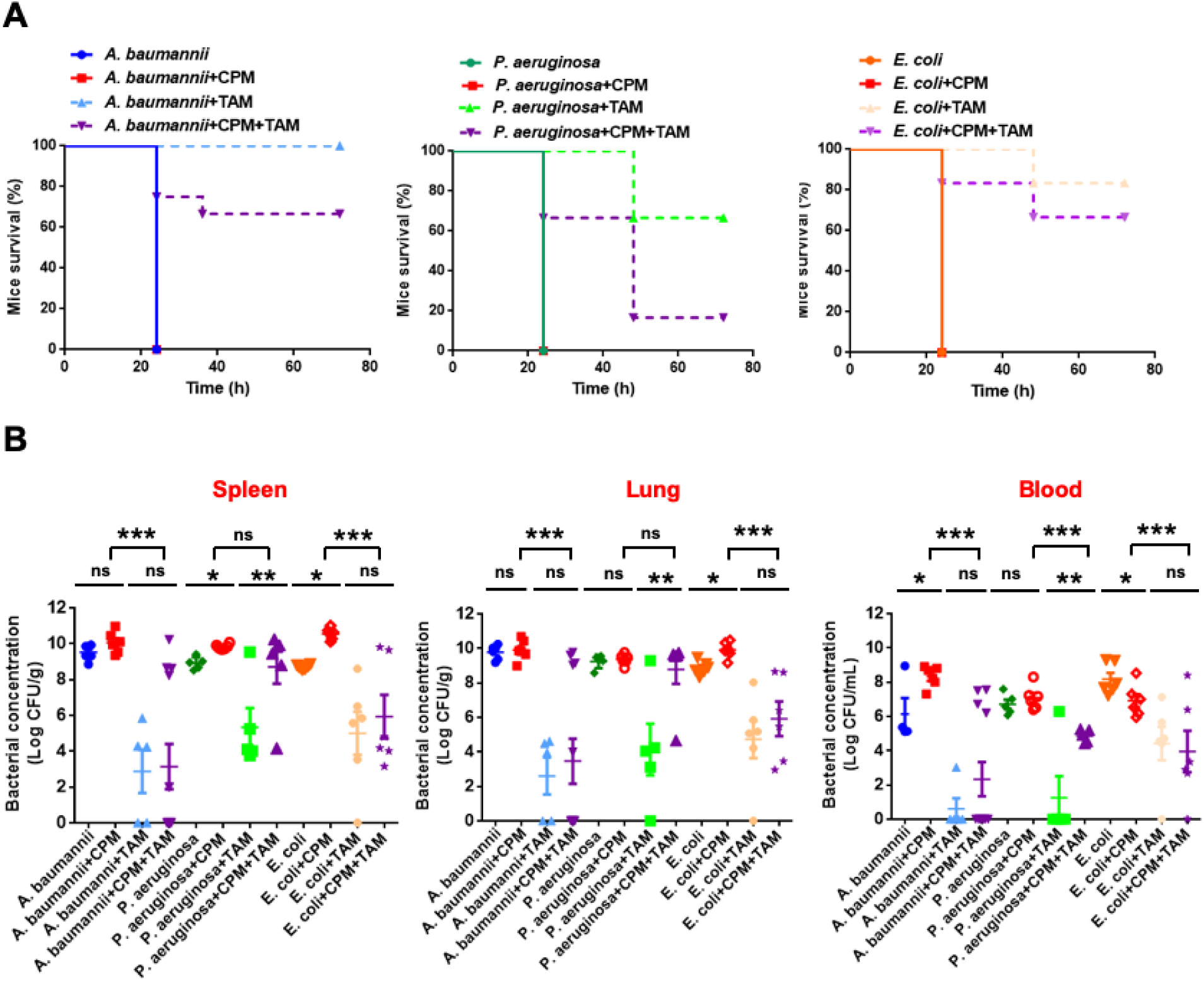
Immunosuppressed mice respond to TAM treatment. (**A**) Mice survival was monitored during 3 days for immunocompetent and neutropenic mice, induced by cyclophosphamide, infected with MLD100 of *A. baumannii* ATCC17978, *P. aeruginosa* PAO1 or *E. coli* ATCC25922 strains treated or not with 3 ip. doses of tamoxifen (80 mg/kg/d, for 3 days). (**B**) Bacterial burden in tissues and blood of immunocompetent and neutropenic mice treated or not with 3 i.p. doses of tamoxifen (80 mg/kg/d, for 3 days), and infected with MLD100 of *A. baumannii* ATCC17978, *P. aeruginosa* PAO1 or *E. coli* ATCC25922 strains. \**P*<0.05: bacteria *vs*. bacteria + CPM, \*\**P*<0.05: bacteria + TAM *vs*. bacteria + CPM + TAM, \*\*\**P*<0.05: bacteria + CPM *vs*. bacteria + CPM + TAM. CPM: cyclophosphamide, TAM: tamoxifen. ns: non-significant.

### Tamoxifen metabolites present antibacterial activity targeting the bacterial membrane

Despite the fact that tamoxifen has no bactericidal activity *in vitro* (MIC > 256 mg/L), we reasoned that the *in vivo* antimicrobial activity of tamoxifen observed in neutropenic mice should result from tamoxifen metabolism in mice organism. Susceptibility assays showed that these tamoxifen metabolites together exhibit MIC_50_ values of 8 and 16 mg/L against 100 and 47 clinical isolates of *A. baumannii* and *E. coli*, respectively (Figure 8A). These data were confirmed by time-kill assays showing that tamoxifen metabolites had excellent bactericidal activity against MDR *A. baumannii* and *E. coli* strains during 8 h of growth (Figure 8B).

**Figure 8.**
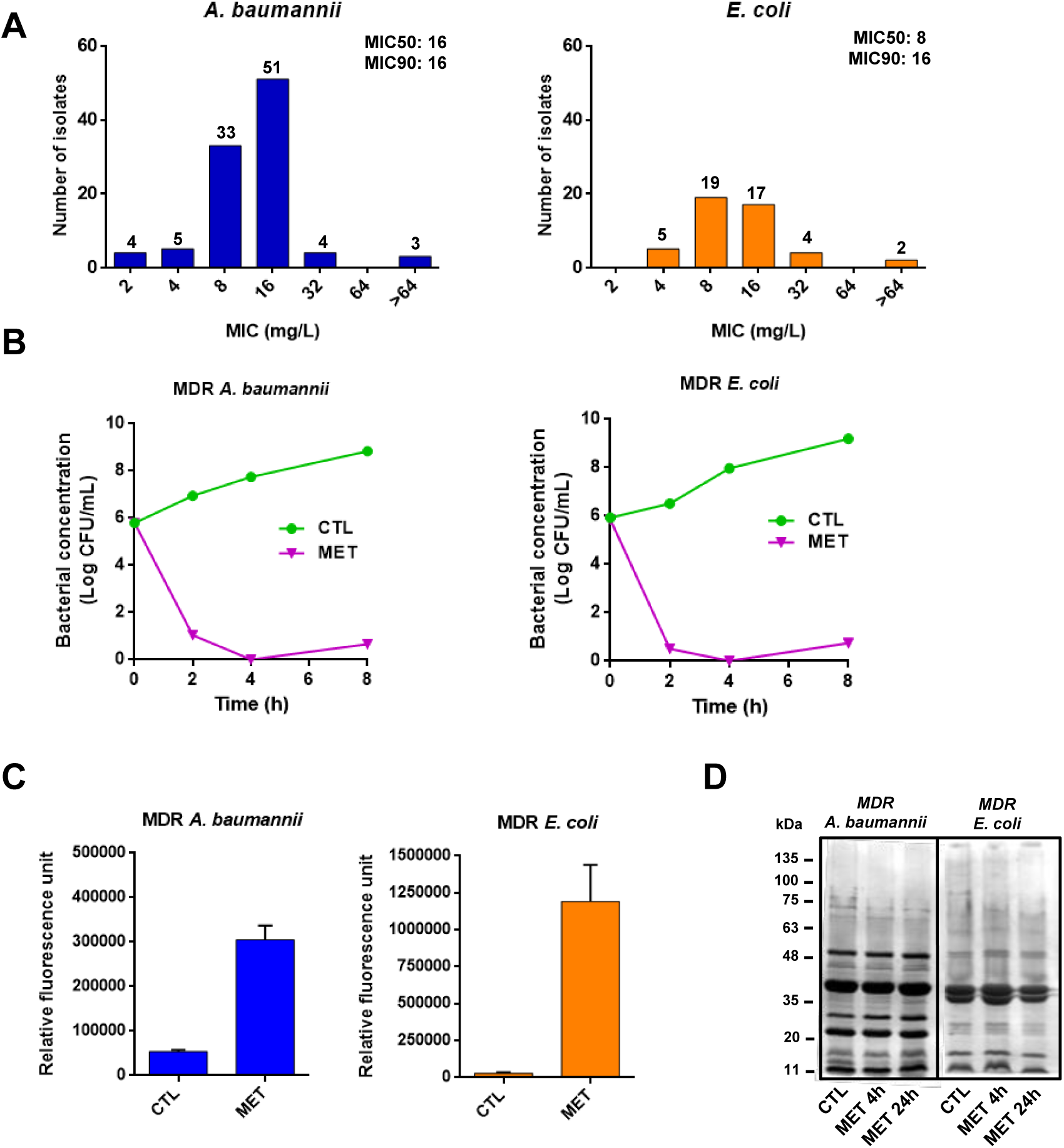
Tamoxifen metabolites present antibacterial activity targeting the bacterial membrane. (**A**) Histogram distribution of MIC for the three tamoxifen metabolites mixture against a collection of *A. baumannii* and *E. coli*. (**B**) Time–kill curves of the MDR *A. baumannii* Ab186 and *E. coli* EcMCR+ strains alone and in the presence of metabolites mixture (4xMIC) for 8 h. (**C**) Tamoxifen metabolites effect on the bacterial permeability. The membrane permeabilization of MDR *A. baumannii* Ab186 and *E. coli* EcMCR^+^ strains in absence and presence of tamoxifen metabolites (2 and 16 mg/L, respectively) incubated for 24 h, was quantified by Typhon Scanner. (**D**) SDS–PAGE of the outer membrane proteins of MDR *A. baumannii* Ab186 and *E. coli* EcMCR^+^ strains with or without tamoxifen metabolites (2 and 16 mg/L, respectively). MET: The three tamoxifen metabolites together. CTL: control.

In order to determine the mode of action of tamoxifen metabolites, we examined their effect on the membrane permeability. Tamoxifen metabolites strongly increased the membrane permeability time-dependent (Figure 8C), without affecting the outer membrane proteins (OMPs) profile (Figure 8D). This suggests that tamoxifen metabolites affect only the integrity of the bacterial cell wall without changing the expression of the OMPs. Determining the specific mechanism of action of tamoxifen metabolites requires further investigation.

## Discussion

The present study provides new data highlighting the antibacterial effect of tamoxifen and its metabolites. Here, we provide the first evidence of an essential role played by tamoxifen in the regulation of immune cells traffic after bacterial infection, in order to reduce the hyperinflammation caused by sepsis, and its antibacterial activity *in vivo* through the generation of active metabolites presenting bactericidal activity against GNB.

This study, as well as previous works (8, 9), showed that the regulation of inflammatory monocytes and neutrophils migration are important in the host defense against bacterial infections. This is consistent with the immune system modulation that improves the bacterial infection clearance (4). Exploiting immunomodulatory drugs, approved by the regulatory agencies for clinical indication different to bacterial infection therapy, has several advantages (30); thus, information of their pharmacological characteristics (toxicity and pharmacokinetics) in preclinical and clinical trials is available. Therefore, the time and the economic costs of the evaluation of these drugs in other therapeutic applications, such as the treatment of bacterial infections, will be reduced (31). Among these immunomodulatory drugs, we found tamoxifen as a promising therapeutic candidate; which has showed antifungal and antibacterial activities against *Mycobacterium tuberculosis* and some Gram-positive bacteria *in vitro* and *in vivo* (17, 18, 25).

Here, we showed that tamoxifen reduces the release of MCP-1 and IL-18, and the phosphorylation of ERK, which contributes to efficient reduction of migration of inflammatory monocytes and neutrophils from bone marrow to blood. Recruitment of both immune cells from bone marrow to blood during systemic infection with GNB is probably mediated by multiple pathways dependent or independent to MCP-1 such as MyD88 and MIP-2 (32-34). MyD88 has been reported to induce MCP-1 release of macrophages after their infection by *Listeria monocytogenes* (32). In contrast, to our knowledge, MIP-2 is not involved in the release of MCP-1 by eukaryotic ells. The presence of pathways independent to MCP-1 has been confirmed in this study in MCP-1 KO mice infected by *A. baumannii, P. aeruginosa* and *E. coli*, which inflammatory monocytes and neutrophils still migrate at lower levels from bone marrow to bloodstream. Previous independent work reported that deletion of MCP-1 in mice did not abolished completely the recruitment of monocytes during the infection by *L. monocytogenes* and this recruitment was diminished by 40-50% (35), suggesting the involvement of MCP-3, another monocyte chemoattractant protein, after binding to CCR2 receptor in the systemic bacterial infection (36). Regarding neutrophils, although is widely accepted that MIP-2 stimulates their migration from bone marrow (37, 38), we demonstrated for the first time that in MCP-1 KO mice the migration of neutrophils from bone marrow to bloodstream after GNB infection was diminished, suggesting the involvement of MCP-1 in this process. This result is consistent with previous observation that MCP-1 regulates the recruitment of neutrophils to the lung after *E. coli* infection (34). Based on these data, MCP-1 plays an important role in the migration of inflammatory monocytes and neutrophils from bone marrow to bloodstream. However, this migration in MCP-1 KO mice infected by GNB and treated with tamoxifen is reduced but not abolished. A possible explanation could be the involvement of other MCP-1-independent pathways regulated by tamoxifen. In this context, further studies are required to decipher the role of these MCP-1-independent pathways in this process.

A consequence of the reduction in monocyte proportions in blood after treatment with tamoxifen would be the reduction of macrophages and dendritic cells in blood and tissues. Although the number of macrophages and neutrophils recruited to the sites of infection in mice treated with tamoxifen would be lower, our *in-vitro* assays suggested that their killing activity against *A. baumannii* and *E. coli* was enhanced by tamoxifen. The inflammatory monocytes are the precursors of a subset of dendritic cells (TipDC), which produce tumor necrosis factor-α (TNF-α) and inducible oxide synthase (iNOS) contributing to the innate defense against *L. monocytogenes* infection (39, 40). In contrast, other study reported that the reduction of proinflammatory monocytes and TipDC during *Trypanosoma brucei* infection diminished their pathogenicity (41). These contradictory results in the effect of monocytes and TipDC recruitment on host survival could be explained by the difference in cellular location of each pathogen, *L. monocytogenes* is intracellular whereas *T. brucei* remains in plasma (*9*). Moreover, it is reported that tamoxifen inhibits *in vitro* the maturation of TipDC, in presence of 17 β-estradiol, which not respond enough to bacterial LPS (42). We suggest that the reduction in the dendritic cells’ proportions joined with their less maturation after tamoxifen treatment produced a reduction in TNF-α and iNOS production, minimizing their deleterious effects in sepsis situation. In our study, we found that mice treatment with tamoxifen reduce the release of proinflammatory cytokines such as TNF-α and IL-6 (data not shown). Accordingly, although we previously pointed that *A. baumannii* could support intracellular lifestyle (43, 44), bacterial species used in our study are viewed as extracellular pathogens and are present in blood. Consequently, it is possible that in our model of study, reduction of monocyte and TipDC frequencies by tamoxifen treatment, and the reduction of proinflammatory cytokines release may play an important role in the therapeutic efficacy of tamoxifen.

It is noteworthy that tamoxifen therapeutic efficacy is not only based in their role regulating the innate immune response, but it is different depending on the type of bacteria. Previous study showed the antibacterial effect of tamoxifen against *Staphylococcus aureus* (25). In the present study, we demonstrate the excellent therapeutic efficacy of tamoxifen against susceptible and MDR *A. baumannii, P. aeruginosa* and *E. coli*, even though this efficacy is slightly lower against *P. aeruginosa*. However, tamoxifen reduced the migration of immune cells from bone marrow to blood in mice infected by these three pathogens at similar levels. A possible explanation could be the involvement of an additional independent immune response mechanism. This hypothesis is in agreement with the results we obtained in neutropenic mice, in which tamoxifen has a therapeutic efficacy against *A. baumannii* and *E. coli*, but not against *P. aeruginosa*. In addition, the three major active metabolites of tamoxifen, 4-hydroxytamoxifen, endoxifen and N-desmethyltamoxifen, as a consequence of its extensive metabolization by cytochrome P450 enzymes (20), present bactericidal activity in monotherapy against *A. baumannii* and *E. coli*, but not against *P. aeruginosa*. These results are consistent with a therapeutic efficacy of tamoxifen depending on antibacterial activity, as addition to the immune response mechanisms. Together, these data indicate that treatment with tamoxifen may be useful for patients with infections by Gram-negative bacilli.

## Materials and Methods

### Reagents

Tamoxifen, N-desmetyltamoxifen, endoxifen and 4-hydroxytamoxifen, porcine mucin, protease inhibitors were obtained from Sigma, Spain. Cyclophosphamide was obtained from Baxter, Spain.

### Bacterial strains

Reference *A. baumannii* ATCC 17978 (45), *P. aeruginosa* PAO1 (46) and *E. coli* ATCC 25922 (47) strains were used. We also used 2 clinical susceptible (Ab9) and multidrug-resistant (MDR) (Ab186) *A. baumannii* from REIPI-GEIH 2010 collection (5), 2 clinical susceptible (Pa39) and MDR (Pa238) *P. aeruginosa* from REIPI-GEIH 2008 collection (48), and 2 clinical susceptible (C1-7-LE) and MDR (EcMCR^+^, carrying *mcr-1* gene) *E. coli* (49, 50). We also used a collection of *A. baumannii* and *E. coli* clinical strains from REIPI-GEIH 2010 collection and Bact-OmpA collection (51, 52).

### Animals

Immunocompetent C57BL/6 female mice (16-18 g) were obtained from the University of Seville facility. MCP-1 KO mice were generated with C57BL/6 background and obtained from Jackson Laboratory, USA. All mice had sanitary status of murine pathogen free and were assessed for genetic authenticity and housed in regulation cages with food and water ad libitum. This study was carried out in strict accordance with the protocol approved by the Committee on the Ethics of Animal Experiments of the University Hospital of Virgen del Rocío, Seville (0704-N-18). All surgery was performed under sodium thiopental anaesthesia and all efforts were made to minimize suffering.

### Immunosuppressed mice

Blood frequencies of monocytes and neutrophils were reduced with cyclophosphamide treatment following the protocol of Zuluaga *et al*. (29). Immunocompetent C57BL/6 female mice were treated with cyclophosphamide at 100 and 150 mg/kg at day 4 and 1, respectively, before the bacterial infection.

### A. baumannii, P. aeruginosa and E. coli peritoneal sepsis models

Murine peritoneal sepsis models with *A. baumannii, P. aeruginosa* or *E. coli* strains were established by ip. inoculation of the bacteria in immunocompetent and neutropenic mice (2). Briefly, 6 mice for each group were inoculated with the minimal bacterial lethal dose 100 (MLD100) of the bacterial suspensions mixed in a 1:1 ratio with a saline solution containing 10% (w/v) porcine mucin. The MLD100 of ATCC 17978, Ab9, Ab186, PAO1, Pa39, Pa238, ATCC 25922, C1-7-LE and EcMCR-1^+^ were 3.2, 5.9, 5.0, 4.9, 3.85, 6.7, 4.7, 2.91 and 6 log CFU/mL, respectively. Mortality was recorded over 3 or 7 days. After the death or sacrifice of the mice at the end of the experimental period, aseptic thoracotomies were performed, and blood samples were obtained by cardiac puncture. The spleen and lungs were aseptically removed and homogenized (Stomacher 80; Tekmar Co., USA) in 2 mL of sterile NaCl 0.9% solution. Ten-fold dilutions of the homogenized spleen, lungs and blood were plated onto Sheep blood agar (Becton Dickinson Microbiology Systems, USA) for quantitative cultures. If no growth was observed after plating the whole residue of the homogenized tissue and blood, a logarithm value corresponding to the limit of detection of the method (1 CFU) is assigned.

### Therapeutic effect of tamoxifen in immunocompetent murine models of peritoneal sepsis

The immunocompetent murine peritoneal sepsis models by *A. baumannii* (ATCC 17978, Ab9 and Ab186), *P. aeruginosa* (PAO1, Pa39 and Pa238), or *E. coli* (ATCC 25922, C1-7-LE and EcMCR-1^+^) strains were established by ip. inoculation of the bacteria in immunocompetent mice. Briefly, 6 animals of each group were infected ip. with 0.5 mL of the MLD100 of each strain mixed 1:1 with 10% porcine mucin. Tamoxifen therapy was administered for 3 days at one safe dose of 80 mg/kg/d before bacterial inoculation. Mice were randomly ascribed to the following groups: 1). controls (without treatment), and 2). Tamoxifen administered at 80 mg/kg/d ip. for 3 days before bacterial inoculation with each strain. Mortality and bacterial loads in tissues and blood were determined as in “*A. baumannii, P. aeruginosa* and *E. coli* peritoneal sepsis models” section.

### Therapeutic effect of tamoxifen in immunosuppressed murine models of peritoneal sepsis

The neutropenic murine peritoneal sepsis models by *A. baumannii* ATCC 17978, *P. aeruginosa* PAO1 or *E. coli* ATCC 25922 strains were established by ip. inoculation of the bacteria in neutropenic mice. Briefly, animals (6 mice for each group) were infected ip. with 0.5 mL of the MLD100 of each strain mixed 1:1 with 10% porcine mucin. Tamoxifen therapy, mortality and bacterial loads in tissues and blood were determined as in “Therapeutic effect of tamoxifen in immunocompetent murine models of peritoneal sepsis” section.

### Flow cytometry

Expanded details of all methods are given in the supplementary material.

### Cytokine assays

Blood samples were collected from periorbital plexuses of mice infected with DML100 of ATCC 17978, PAO1 and ATCC 25922 and pretreated or not with tamoxifen as previously described (11). Serum levels of murine MCP-1, IL-6, IL-18 and TNF-α were collected 6 and 24 h post-bacterial infection without or with tamoxifen treatment. MCP-1, IL-6, IL-18 and TNF-α levels were determined by ELISA kit (ThermoFisher, for MCP-1) and (Affymetrix eBioscience, for IL-6, IL-18 and TNF-α) in accordance with the manufacturer’s instructions. Furthermore, extracellular medium of RAW 264.7 macrophages cells infected with 8 log CFU/mL of ATCC 17978, PAO1 and ATCC 25922, and pre-incubated or not with 2.5 mg/L tamoxifen for 24 h previous was collected to determine the MCP-1 levels.

### Cell culture and infection

Expanded details of all methods are given in the supplementary material.

### Western blot immunoblotting

Expanded details of all methods are given in the supplementary material.

### Macrophages adhesion assay

RAW 264.7 cells were pretreated with 2.5 mg/L tamoxifen for 2, 6 and 24 h; and infected with ATCC 17978, PAO1 and ATCC 25922 strains (MOI 1:100) for 2 h with 5% CO2 at 37°C. Subsequently, infected RAW 264.7 macrophages cells were washed five times with prewarmed PBS and lysed with 0.5 % Triton X-100. Diluted lysates were plated onto Sheep blood agar and incubated at 37 °C for 24 h for enumeration of developed colonies and then the determination of the number of bacteria that attached and invaded RAW 264.7 cells. Alternatively, we determined the concentration of the extracellular medium bacteria by plating diluted extracellular medium onto Sheep blood agar.

HL-60 neutrophils were pretreated with 2.5 mg/L for 2 and 6 h; and infected with ATCC 17978, PAO1 and ATCC 25922 strains (MOI 1:100) for 2 h with 5% CO2 at 37°C. Subsequently, HL-60 neutrophils were washed five times with prewarmed PBS by centrifugation and lysed with 0.5 % Triton X-100. Diluted lysates were plated onto Sheep blood agar and incubated at 37 °C for 24 h for enumeration of developed colonies and then the determination of the number of bacteria. The neutrophil killing index was calculated according to the formula: [(CFU in the absence of neutrophils - CFU in the presence of neutrophils)/ CFU in the absence of neutrophils] × 100 (53).

### In vitro susceptibility testing and time-kill experiments

The MICs of N-desmetyltamoxifen, endoxifen and 4-hydroxytamoxifen and the mixture of the 3 tamoxifen metabolites against *A. baumannii* and *E. coli* clinical strains were determined by microdilution assay in two independent experiments, in accordance with CLSI guideline (54).

Time-kill kinetic assays of the Ab9 and EcMCR^+^ strains were conducted in Moeller Hinton Broth in the presence of the mixture of the 3 tamoxifen metabolites at 4xMIC, and were performed in duplicate as previously described (54). Drug-free broth was evaluated in parallel as a control and cultures were incubated at 37°C. Viable counts were determined by serial dilution at 0, 2, 4 and 8 h after adding the 3 tamoxifen metabolites, and plating 100 μL of control, test cultures or dilutions at the indicated times onto sheep blood agar plates. Plates were incubated for 24 h and, after colony counts, the log10 of viable cells (CFU/mL) was determined.

### Analysis of outer membrane proteins (OMPs) by SDS-PAGE

Bacterial cells of MDR *A. baumannii* and MDR *E. coli* were grown in LB broth to the logarithmic phase, incubated with 2 and 16 mg/L of tamoxifen metabolites mixture, respectively, for 4 or 24 h and lysed by sonication. OMPs were extracted with sodium lauroylsarcosinate (Sigma, Spain) and recovered by ultracentrifugation as described previously (43). The OMP profiles were determined by sodium dodecyl sulfate polyacrylamide gel electrophoresis (SDS-PAGE) using 10% SDS gels and 6 μg protein of OMPs, followed by Simply Blue SafeStain gel (Invitrogen, Spain).

### Membrane permeability assays

Bacterial suspensions (adjusted to O.D_600_ = 0.2) of MDR *A. baumannii* and MDR *E. coli* were placed into a 96-well plate, incubated with 2 and 16 mg/L of tamoxifen metabolites mixture, respectively, and mixed in a solution of PBS containing Ethidium Homodimer-1 (EthD-1) (1:500) (Invitrogen, USA). After 10 min of incubation, fluorescence was monitored during 160 min using Thyphoon FLA 9000 laser scanner (GE Healthcare Life Sciences, USA) and quantified by ImageQuant TL software (GE Healthcare Life Sciences, USA). Bacterial counts were obtained at the beginning and at the end of the experiment to ensure that metabolites mixture was not presenting bactericidal activity against *A. baumannii* and *E. coli* strains.

### Statistical analysis

Group data are presented as means ± standard errors of the means (SEM). For *in vitro* studies, the Student t test was used to determine differences between means. Differences in bacterial spleen, lung and blood concentrations (mean ± SEM log_10_ CFU per g or mL) were assessed by analysis of variance (ANOVA) and post-hoc Dunnett’s and Tukey’s tests. Differences in mortality (%) between groups were compared by use of the χ2 test. *P* values of <0.05 were considered significant. The SPSS (version 21.0; SPSS Inc.) statistical package was used.

## Acknowledgments

We thank Dr. José Manuel Rodríguez Martinez for the kind gift of the EcMCR+ strain.

## Author contributions

R.T., J.P., Y.S. conceived the study and designed the experiments. A.M.C., R.A.A, R.T, performed experiments and interpreted data. J.P. and Y.S. wrote the manuscript with the input of all the other authors.

